# Pioneer transcription factors in chromatin remodeling: the kinetic proofreading view

**DOI:** 10.1101/2020.01.24.916908

**Authors:** Helmut Schiessel, Ralf Blossey

## Abstract

Pioneer transcription factors are a recently defined class of transcription factors which can bind directly to nucleosomal DNA; they play a key role in gene activation in certain pathways. Here we quantify their role in the initiation of nucleosome displacement within the kinetic proofreading scenario of chromatin remodeling. The model allows to perform remodeling efficiency comparisons for scenarios involving different types of transcription factors and remodelers as a function of their binding and unbinding rates and concentrations. Our results demonstrate a novel way to fine-tune the specificity of processes that modify the chromatin structure in transcriptional initiation.

## Introduction

The kinetic proofreading scenario of chromatin remodeling was originally formulated in order to provide a conceptual and quantitative answer to the simple question: how does a remodeler decide on the right nucleosome to remodel at a given time? Our theory gave a tentative answer based on the idea that specific histone-tail states and the ATP-consumption of remodeling enzymes act together to bias the action of remodelers towards specific nucleosomes [1]. This idea is much in line with the original concept of kinetic proofreading developed by Hopfield [2] and Ninio [3], with the key difference that kinetic proofreading in the chromatin remodeling context refers to the kinetic selection of a substrate as a dynamic intermediate—the nucleosome to be displaced—rather than the proofreading of a product, as in the classic proofreading example of mRNA translation into a protein.

The kinetic proofreading scenario of chromatin remodeling found a first experimental verification through the work of G. J. Narlikar [4, 5], who developed essentially the same idea based on the experiments with ISWI-type remodelers performed in her group [6]. Recently, important new experimental insights were gained with high-throughput remodeling assays [7], this time for a larger group of members from the ISWI-family of remodelers. We recently reanalyzed these data, for which no model interpretation was offered in [7], in the context of our kinetic proofreading scenario [8].

In Ref. [8], we have presented a list of further examples of chromatin remodeler-nucleosome interactions, which are useful for putting our scenario to experimental scrutiny. This test is most relevant for members of other remodeler families which have different histone-tail specificities and also other ATP-characteristics than IWSI-type remodelers. Another one of these examples refers to the role of transcription factors in the recruitment of chromatin remodelers. Already in our 2008 paper we were aware of the fact that the specific recognition of nucleosome based histone-tail modifications alone might not be the only relevant mechanism, depending on the cellular context. We therefore had included transcription factors (TFs) (see Figure 1 of [1]) as additional contributors of specificity, but we did not have enough details at hand on how these factors might specifically intervene. In this paper we return to this issue building on very recent experimental insights.

**FIG. 1.**
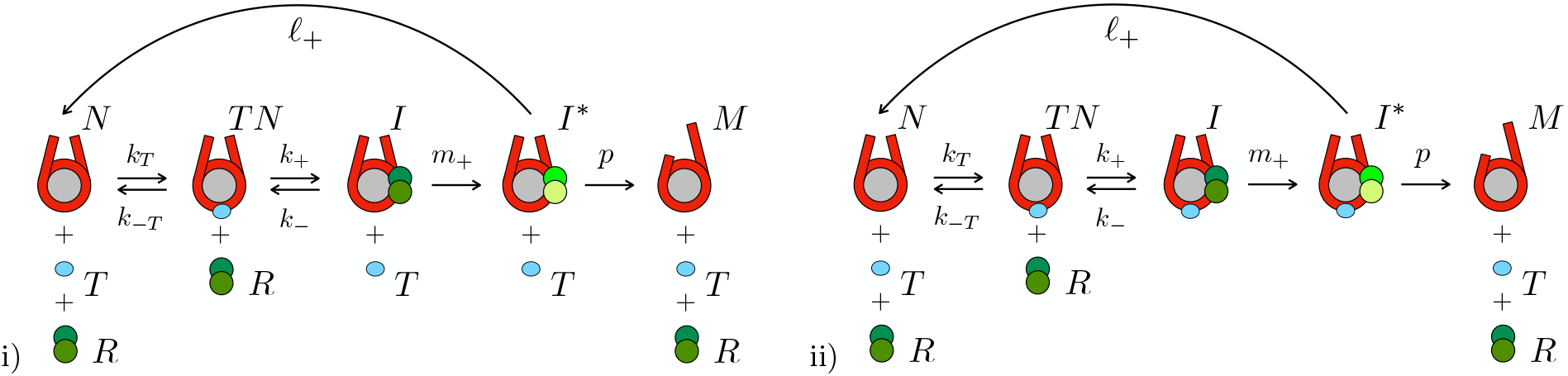
Kinetic proofreading scheme for chromatin remodeling with nucleosome (*N*), pioneer transcription factor (*T*), remodeler (*R*). Two scenarii are shown: i) early pTF unbinding; ii) late pTF unbinding. In case i), *I* denotes the remodeler-nucleosome complex and *I*\* the activated complex; *M* is the mobile nucleosome. In case ii) *I* and *I*\* stand for the corresponding remodeler-nucleosome-pTF complexes. The translocation step with rate *p* is shown to ultimately lead to remodeler dissociation; this step is in general processive.

Transcription factors have been shown to intervene in several different ways in transcriptional processes in chromatin; quite generally they have been shown to compete with nucleosomes for binding to DNA [9–12] and they also interact with chromatin remodelers [13]. These interactions of transcription factors with the nucleosomes are essentially controlled by direct and indirect sequence specificities and otherwise dominated by steric exclusion effects and therefore have probably only little or no specific regulatory relevance for the initial recruitment of chromatin remodelers; they might, however, be important for the maintenance of transcriptionally active states.

Recently, however, a different class of transcription factors has come into view, which potentially plays decisive roles for early regulatory events involving the recruitment of chromatin remodelers. These so-called pioneer transcriptions factors (pTFs) engage with nucleosomes present in condensed nucleosomal arrays, and can bind directly to nucleosomes [15–19]. A recent review details the different structural motifs through which the transcription factors can bind to nucleosomal DNA [20]. One exemplary pTF is FoxA, whose DNA-binding domain has sequence similarity with histone H1—this example immediately makes clear that pTFs have DNA-binding domains that exploit the characteristics of nucleosomal DNA [15, 21]. More recently, the exemplary case of BZLF1 has been scrutinized in detail [22]; this pTF is involved in the activation of the Epstein-Barr virus (EBV) in the human genome, where it induces a lytic behaviour from a latent state—much in analogy to the famous λ-phage in *E. coli*—only now in the context of a human disease and, on the molecular level, in the context of chromatin. Another very recent work details the role of the pTF Rap1 in a combined *in-vitro* and *in-vivo* study in both binding to nucleosomes and recruiting the chromatin remodeler RSC [23].

What can thus be taken as established from these papers is that pTFs bind to nucleosomal DNA and interact with specific chromatin remodelers to initiate the opening of nucleosomal arrays. In the following section we will show how these two key properties of pTFs can be integrated into our remodeling model and to what predictions these assumptions lead for remodeler discrimination.

## Model

The role pTFs play in the regulation of chromatin remodeling can be understood from a modification of the rate-equation based kinetic proofreading model [8]. The model considers four molecular partners: the nucleosome *N*, the remodeler *R*, and remodeler nucleosome complexes *I* and *I*\*, whereby the latter refers to the activated state upon ATP-consumption. Here, we extend this model by introducing the pTF *T*. The scheme we propose for this extended system is given by the following four reactions (see also Fig. 1 i)):

a. binding of the pTF to the nucleosome:

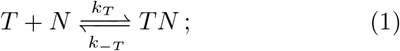
b. binding of the remodeler to the nucleosome carrying the pTF, formation of the remodeler-nucleosome complex and unbinding of the pTF,

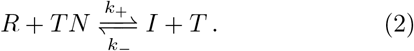 This reaction requires some additional interpretation since part of the back reaction with rate *k*_−_ is already covered by the reaction with rate *k_T_* in (1). Reaction (2), as it stands, should be understood as consisting of the two reactions

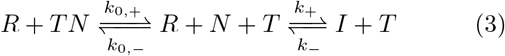

whereby the first reaction should be read as a fast unbinding of the pTF *T* before the remodeler *R* binds to the nucleosome *N* and forms the remodeler-nucleosome complex *I*. The second slow reaction then essentially corresponds to the complex formation. Then, we have as the next reaction
c. activation of the remodeler-nucleosome complex:

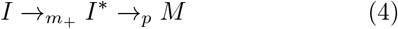

where the activated intermediate *I*\* is translocated and hence becomes a mobile nucleosome, which we call M. And, finally, we have
d. unbinding of the remodeler after termination of the remodeling processes via

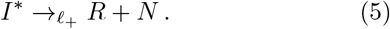 These reactions can be easily expressed in the form of first-order differential equations. In the absence of the pTF, one has two equations for the remodeler-nucleosome complexes [*I*] and [*I*\*] [8]; one now has three:

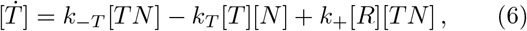

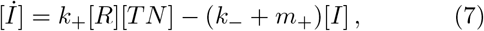

and

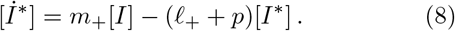

The efficiency of the remodeling process can be quantified by considering a stationary process and computing the quantity

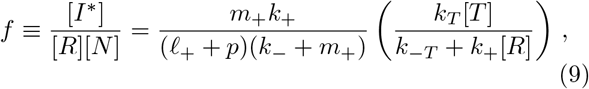

which expresses the concentration ratio of active nucleosome-remodeler complexes to their independent components; we call it an efficiency coefficient for brevity. We see that we can write this expression as the product

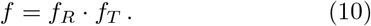

This efficiency coefficient thus factorizes, whereby the first factor *f_R_* is the remodeler efficiency one encounters for chromatin remodeling in the absence of pTFs [8], while the second factor

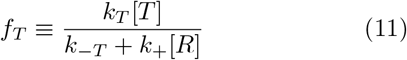

arises due to the presence of pTF-binding.

The liberation of the transcription factor *T* from the nucleosome, assumed in Eq. (2) need not necessarily be the case, see e.g. [23]. We can also have the case in which the pTF stays on the nucleosome together with the remodeler-nucleosome complex and only detaches upon mobilization of the complex and its transport along DNA. In this case, while reaction (1) stays unchanged, the following reactions are modified. Reaction b) is replaced by b2), the binding of the remodeler to the nucleosome carrying the pTF without the unbinding of the latter. *I*then stands for the remodeler-nucleosome-pTF complex and fulfills the equation

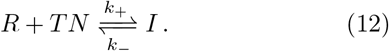

Further, c2) is the activation of the remodeler-nucleosome-pTF complex is now modified by the dissociation of the pTF

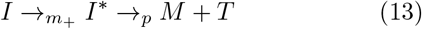

where the activated intermediate *I*\* is translocated and hence becomes a mobile nucleosome, which we call M. And, finally, we have the unchanged reaction d), the unbinding of the remodeler after termination of the remodeling processes.

The new reaction scheme a), b2), c2), d) leads to only one change, namely in the equation for [*T*], Eq. (6), in which the last term is modified. It reads as

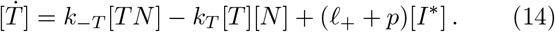

The remodeling efficiency *f* in this case is modified as the rate *k*_+_ in the factor *f_T_* is normalized with a factor α, *i.e*.

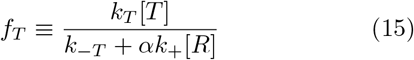

where *α* is given by

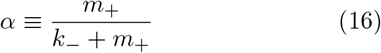

which incidentally also is part of the factor *f_R_*. Given that for a successful remodeling process one has *k*_−_ ≪ *m*_+_, the factor *α* in general does not deviate much from one and can therefore be safely neglected in the following discussion.

## Discussion of recruitment efficiencies

The new factor (11) shows that the remodeler efficiency *f* in the presence of pTFs is not just rate-dependent, as in the case of remodelers alone [8], but also depends on pTF and remodeler concentrations. One easily distinguishes two limits from the denominator of (11):

i. if *k_−T_* ≪ *k*_+_[*R*], one has

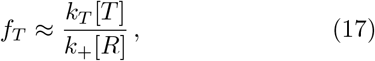

while in the opposite case:
ii. *k_−T_* » *k*_+_[*R*]

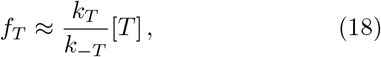

*i.e*., the remodeler efficiency is modified by an equilibrium binding factor due to the involvement of the pTF.

In the next step we can now discuss the comparison of remodeler efficiencies for different scenarios in order to find out about which scenario will be favored. The quantity to compute then is the ratio *F* = *f*_1_/*f*_2_ of two efficiencies *f_i_* which can now be identified with different situations; the most relevant being the distinction between two different remodelers *R*_1_ and *R*_2_. It is given by

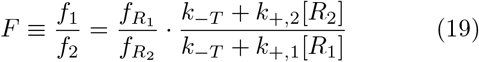

which clearly shows a dependence on the concentration of the two competing remodelers. The other limiting situation arises when one considers the discrimination of one remodeler with respect to two different pTFs, in which case one has

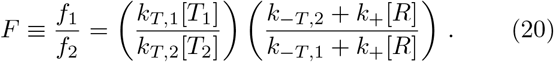

## Quantitative estimates

We will now provide some order-of-magnitude quantitative estimate of how the presence of the pTF in the reaction kinetics alters remodeling efficiencies. For these we use results from highly quantitative work on transcription factor binding [24] in combination with results from the recent experiments on the pTF Rap1 [23].

We consider the factor (11) by looking at its slightly reorganized inverse

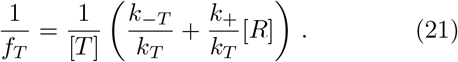

We take from [24] the value for the binding rates as *k_T_* ≈ 0.1 s^−1^ nM^−1^, see caption of Figure 4 in that paper and the Supplementary Material. We assume this value holds approximately also for the chromatin remodeler binding rate *k*_+_. The first term in (21) is the dissociation constant of the pTF; here, the case of Rap1 on nucleosomes positioned on the RPL30 promoter is particularly illuminating: for the two different positions of pTF-binding at nucleosomal SHL 4.5 and SHL 6.5 the dissociation constant *K_D_* = *k_−T_*/*k_T_* was found to have values of 10 nM and 30 nM, respectively [23]. We thus see we have *k_−T_* » *k*_+_ [*R*] provided we consider remodeler concentrations in the low nM ranges; this leads us to case ii) discussed in the previous section. We see that in this case, the modification of the remodeling efficiency depends in fact on the specific binding position of the pTF to the nucleosome. The experiments in [23] were designed to detect clear differences in remodeling in the case of the presence of at least one binding motif, or the absence of both.

## Discussion and Conclusions

In this paper we have proposed to explain the action of pioneer transcription factors (pTFs), as recently described in experiments for the first time, in the context of the kinetic proofreading scenario of chromatin remodeling. pTFs are key factors to induce an open chromatin state from condensed arrays and therefore play a key role in gene activation. The inclusion of pTFs in the kinetic proofreading scenario shows that pTFs can act as enhancers of remodeler recruitment, which illustrates very clearly the original vision of the model in that an equilibrium binding step - here leading to a nucleosome-bound transcription factor - biases the remodeling step towards the ‘right nucleosome to remodel at a given time’.

The new equations (19) and (20) for the discrimination ratio are particularly illuminating in this respect as they manifest the active role played by the involvement of different remodelers and transcription factors in gene activation. In contrast to the case studied before, in which discrimination is governed by internal nucleosomal states, notably the spectrum of histone tails, the new discrimination ratio show dependencies both on reaction rates and on the respective *concentrations* of these molecules.

Within the kinetic proofreading scheme this constitutes a novel control mechanism of chromatin remodeling regulation which depends directly *on the expression levels of pTFs and remodelers* rather than only on the kinetics of their interaction.

Via eqs. (19) and (20) our model allows us to calculate simple expressions for remodeler discrimination due to the presence of the pioneer transcription factors which, depending on the magnitude of the involved rates and concentrations of both pTFs and remodelers, describe different modes of discrimination between nucleosomes for remodeling. The model thus yields new testable predictions for experiments on chromatin remodeling involving different pTFs and remodelers both *in vitro* and *in vivo*; it is also easily adaptable to additional situations encountered in experiment. *In vivo*, following the exemplary study of the combined action of remodelers in nucleosome positioning [25], such experiments can yield important insights into molecular concentration effects in chromatin remodeling.

In summary, our theoretical results - originally motivated by a speculative idea about chromatin regulation - find a new biologically relevant application to the involvement of pioneer transcription factors in gene activation and therefore reach beyond the recent application of our scenario to high-throughput experiments on members of the ISWI-family of remodelers [8]; the development of this model variant became possible only due to experimental insights obtained in the last months. Based on our extended model we propose a new way in which pTF-nucleosome-remodeler interactions can be fine-tuned in the nucleus not only by affecting binding and reaction rates, but by the expression levels of pTFs and remodelers. The qualitative and quantitative predictions of the model can be experimentally tested both in *in vivo* and *in vitro* settings and hence provide a new path to the validation of the kinetic proofreading scenario of chromatin remodeling.

## Acknowledgement

RB thanks John F. Marko for a discussion on his work on transcription factor binding.

## References

[1] R. Blossey and H. Schiessel, HFSP J. 2 167 (2008).

[2] J. J. Hopfield, Proc. Natl. Acad. Sci. USA 71, 4135 (1974).

[3] J. Ninio, Biochimie 57, 587 (1975).

[4] G. J. Narlikar, Curr. Opin. Chem. Biol. 14, 660 (2010).

[5] R. Blossey and H. Schiessel, Biophys. J. 101, L30 (2011).

[6] T. R. Blosser, J. G. Yang, M. D. Stone, G. J. Narlikar, and X. Zhuang, Nature 462, 1022 (2009).

[7] G. P. Dann, G. P. Liszczak, J. D. Bagert, M. M. Muöller, U. T. T. Nguyen, F. Wojcik, Z. Z. Brown, J. Bos, T. Panchenko, R. Pihl, S. B. Pollock, K. L. Diehl, C. D. Allis, and T. W. Muir, Nature 548, 607 (2017).

[8] R. Blossey and H. Schiessel, Phys. Rev. E 99, 060401(R) (2019).

[9] M. Li, A. Hada, P. Sen, L. Olufemi, M. A. Hall, B. Y. Smith, S. Forth, J. N. McKnight, A. Patel, G. D. Bowman, B. Bartholomew, and M. D. Wang, eLife 4, e06249 (2015).

[10] S. R. Grossman, J. Engreitz, J. P. Ray, T. H. Nguyen, N. Hacohen, and E. S. Lander, Proc. Natl. Acad. Sci. USA 115, E7222 (2018).

[11] S. Rudnizky, H. Khamis, O. Malik, P. Melamed, and A. Kaplan, Proc. Natl. Acad. Sci. USA 116, 12161 (2019).

[12] S. Brahma and S. Henikoff, Trends in Biochem. Sci. 45, 13 (2020).

[13] D. Barisic, M. B. Stadler, M. Iurlaro, and D. Schübeler, Nature 569, 136 (2019).

[14] J. Ren, R. Finney, K. Ni, M. Cam, and K. Muegge, Epigenetics 14, 277 (2019).

[15] L. A. Cirillo, F. R. Lin, I. Cuesta, D. Friedman, M. Jarnik, and K. S. Zaret, Molecular Cell 9, 279 (2002).

[16] E. E. Swinstead, V. Paakinaho, D. M. Presman, and G. L. Hager, Bioessays 38, 1150 (2016).

[17] A. Mayran and J. Drouin, J. Biol. Chem. 293, 13795 (2018).

[18] E. T. Friman, C. Deluz, A. C. A. Meireles-Filho, S. Govindan, V. Gardeux, B. Deplancke, and D. M. Suter, eLife 8 e50087 (2019).

[19] M. M. Makowski, G. Gaullier, and K. Luger, J. Biosci. 45:13 (2020)

[20] M. F. Garcia, C. D. Moore, K. N. Schulz, O. Alberto, G. Donague, M. M. Harrison, H. Zhu, and K. S. Zaret, Molecular Cell 75, 921 (2019).

[21] F. M. Cernilogar, S. Hasenöder, Z. Wang, K. Scheibner, I. Burtscher, M. Sterr, P. Smialowski, S. Groh, I. M. Evenroed, G. D. Gilfillan, H. Lickert, and G. Schotta, Nucl. Acids. Res. 47, 9069 (2019).

[22] M. Schaeffner, P. Mrozek-Gorska, A. Buschle, A. Woellmer, T. Tagawa, F. M. Cernilogar, G. Schotta, N. Krietenstein, C. Lieleg, P. Korber, and W. Hammerschmidt, Life Science Alliance 2, e201800108 (2019).

[23] M. Mivelaz, A.-M. Cao, S. Kubik, S. Zencir, R. Hovius, I. Boichenko, A. M. Stachowicz, C. F. Kurat, D. Shore, and B. Fierz, Mol. Cell. 77, 1–13 (2020)

[24] R. I. Kamar, E. J. Banigan, A. Erbas, R. D. Giuntoli, M. Olvera de la Cruz, R. C. Johnson, and J. F. Marko, Proc. Natl. Acad. Sci. USA 114, E3251 (2017).

[25] T. Gkikopoulos, P. Schofield, V. Singh, M. Pinskaya, J. Mellor, M. Smolle, J.L. Workman, G.J. Barton, and T. Owen-Hughes. Science 333 1758–1760 (2011).

